# Placental Nanoparticle-mediated IGF1 Gene Therapy Corrects Fetal Growth Restriction in a Guinea Pig Model

**DOI:** 10.1101/2024.04.05.587765

**Authors:** Baylea N Davenport, Rebecca L Wilson, Alyssa A Williams, Helen N Jones

## Abstract

Fetal growth restriction (FGR) caused by placental insufficiency is a major contributor to neonatal morbidity and mortality. There is currently no in utero treatment for placental insufficiency or FGR. The placenta serves as the vital communication, supply, exchange, and defense organ for the developing fetus and offers an excellent opportunity for therapeutic interventions. Here we show efficacy of repeated treatments of trophoblast-specific human *insulin-like 1 growth factor* (*IGF1*) gene therapy delivered in a non-viral, polymer nanoparticle to the placenta for the treatment of FGR. Using a guinea pig maternal nutrient restriction model (70% food intake) of FGR, nanoparticle-mediated *IGF1* treatment was delivered to the placenta via ultrasound guidance across the second half of pregnancy, after establishment of FGR. This treatment resulted in correction of fetal weight in MNR + *IGF1* animals compared to sham treated controls on an ad libitum diet, increased fetal blood glucose and decreased fetal blood cortisol levels compared to sham treated MNR, and showed no negative maternal side-effects. Overall, we show a therapy capable of positively impacting the entire pregnancy environment: maternal, placental, and fetal. This combined with our previous studies using this therapy at mid pregnancy in the guinea pig and in two different mouse model and three different human in vitro/ex vivo models, demonstrate the plausibility of this therapy for future human translation. Our overall goal is to improve health outcomes of neonates and decrease numerous morbidities associated with the developmental origins of disease.

## INTRODUCTION

10% of humans worldwide are born too small and too early ^1^. This equates to 32 million fetal growth restricted (FGR) newborns including over 2.5 million stillbirths and 15 million preterm deliveries ^2-5^. When not victim to stillbirth, FGR neonates are at increased risk of developing complications and morbidities associated with developmental origins of health and disease (DOHaD), which links FGR-associated fetal programming to increased risk of childhood/adult morbidities such as developmental delays/cognitive deficits, cardiovascular disease, and obesity ^6, 7^. There is currently no treatment for FGR or placental insufficiency, defined by the placenta failing to transfer enough nutrients and oxygen for proper fetal growth. The only intervention is early delivery which leads to extended NICU stays. While medicine has made great progress with neonatal care, our ultimate goal is to achieve normal gestational length with upward fetal growth trajectories as long as possible for optimal growth since NICU admissions still have significant negative physical and mental impacts on the infant and parents ^8, 9^ While there are many intrinsic idiopathic causes of FGR stemming from placenta malformation/function or fetal genetic anomalies, there are also numerous extrinsic causes. Extrinsic causes include maternal stress and maternal comorbidities such as diabetes, nutrient deprivation, or drug/alcohol use. However, the commonality between these intrinsic and extrinsic causes is failure of the placenta (whether due to maldevelopment or malfunction) to provide enough nutrients and oxygen to the fetus for appropriate growth ^10-12^. The placenta serves as the vascular, endocrine, and circulatory organ responsible for communication between parent and fetus and offers an excellent opportunity for therapeutic interventions: targeting the origin of the disorder, treating an organ that is discarded after pregnancy, and not directly intervening with the fetus ^13^. By correcting fetal growth restriction, the number of stillbirths, premature neonates, and adult comorbidities could be dramatically reduced for an overall healthier human population.

The insulin like growth factor (IGF) signaling axis is a master regulator of placental development and function, regulating cell proliferation/differentiation, vasculature structure, nutrient transport, and hormone production. IGF1 is downregulated in human FGR cases and vital for placental function and fetal growth in transgenic mouse studies ^14, 15^. Many other growth factors including vascular endothelial growth factor (VEGF), placental growth factor (PLGF), and epidermal growth factor (EGF) have been investigated for the correction of FGR, but only IGF1 acts to regulate all of these processes ^16-18^. Unlike the other growth factors mentioned which are temporally produced in the placenta/maternal-fetal interface, IGF1 is produced and necessary throughout the entirety of gestation. Previous studies aiming to increase fetal growth with IGF1 have used viral delivery, IGF1 protein, or delivered through either the maternal blood supply or amniotic fluid ^19-21^. However, we aim to correct placental insufficiency and FGR through non-viral delivery of DNA directly targeting the placental trophoblast for potential clinical use.

In contrast to previous studies using IGF1 protein, we have developed a non-viral, biodegradable copolymer complexed with a plasmid containing the *IGF1* gene under control of a trophoblast-specific promoter to produce a nanoparticle capable of delivering placenta specific, transient (does not integrate into the genome) transgene expression ^19, 20, 22-28^. Thus far in our nanoparticle therapy studies we have successfully delivered plasmids *ex vivo* and *in vitro* to human syncytiotrophoblast and *in vivo* to mouse, guinea pig, and non-human primate models and shown physiochemical properties, cellular safety, and efficiency in both mother and fetus ^24, 25, 29^. We have also shown that administration of this gene therapy to the placenta in a normal pregnancy environment results in homeostatic responses within the placenta maintaining normal fetal growth and no placentomegaly ^23, 29^. Previous studies using the guinea pig model of FGR from our lab have focused on short term (one *IGF1* nanoparticle treatment with samples collected 5 days later) effects of our nanoparticle-mediated *IGF1* on the placenta at mid-pregnancy and showed positive improvements to the placenta that could improve fetal growth ^24^. Building on from this, this study employed multiple *IGF1* treatments across later gestation to assess longer term impact on placental function, fetal weight and maternal environment.

In this study we used the well-established, non-invasive maternal nutrient restriction model in the guinea pig, a model of maternal stress which establishes placental insufficiency ^30, 31^. Guinea pigs are an excellent model to study placental and fetal development. While mouse models offer genetic tools and many established techniques, the mouse does not offer the best model for placental or fetal development for human translation, as they do not have invasive haemomonochorial placentas and many organ systems finish developing postnatally ^32, 33^. Non-human primate studies are currently underway to establish dosing and safety in a pre-clinical model but do not provide a system to model FGR across late pregnancy reflecting the human ^29^. Guinea pigs, however, provide a model with similar developmental milestones of fetal and placental development compared to humans including similar hormonal profiles, similar placental structure, and similar fetal and neonatal organ development timelines ^32-34^. Here we aimed to identify if repeated delivery o*f* nanoparticle-mediated *IGF1* treatment can correct fetal growth by restoring aberrant placental physiology and signaling and determine impact on the maternal environment. Key aspects of this study include the establishment of the growth restriction prior to treatment at clinically relevant timepoints for human pregnancy/diagnosis and allowing treatment to begin after this diagnosis.

## METHODS

### Nanoparticle Formation

Nanoparticles were formed by complexing a non-viral PHPMA115-b-PDMEAMA115 co-polymer with plasmids containing the *IGF1* gene under the control of a trophoblast-specific promoter, *CYP19A1* (50 µg plasmid in a 200 µL volume) at room temperature. Detailed methods of copolymer synthesis and nanoparticle formation can be found in Wilson et al., 2022 ^24^.

### Animal Husbandry and Nanoparticle Delivery

Animal care and usage was approved by the Institutional Animal Care and Usage Committee at the University of Florida (Protocol #202011236). Power analysis based on data from our previous study ^24^ determined that 6 female guinea pigs per group would be needed to detect a difference (effect size = 1.4, or larger than 1 standard deviation difference) of P≤0.05 with 80% power for outcomes including reduced fetal weight and increased fetal glucose concentrations. Female Dunkin-Hartley guinea pigs (Dams) were purchased (Charles River Laboratories, Wilmington, MA) at 500–550 g (approximately 8-9 weeks of age) and housed in an environmentally controlled room (22°C/72°F and 50% humidity) under a 12 h light-dark cycle. Upon arrival, food (LabDiet diet 5025: 27% protein, 13.5% fat, and 60% carbohydrate as % of energy) and water were provided ad libitum. After a 2-week acclimation period, dams were weighed, ranked from heaviest to lightest and systematically assigned to either ad libitum diet (termed Control: n = 6) or maternal nutrient restriction (MNR) diet (n=12) consisting of a 70% food intake diet (per kilogram body weight of control group) from 4 weeks prior to mating through mid-pregnancy (GD 35), then increased to 90% food intake until end of term to maintain pregnancies. Timed matings, pregnancy confirmation ultrasounds, and ultrasound-guided intra-placental nanoparticle treatments were performed as previously described ^24^. Briefly, dams were anesthetized, and nanoparticle was delivered to the placenta via ultrasound-guided intra-placental injection, 8 days apart at GD36, 44, and 52 (±3). Only one placenta per litter was injected with either nanoparticle-mediated *IGF1* (n=6 MNR + *IGF1*) or a non-expressing sham nanoparticle (n=12; 6 Control and 6 MNR) (Supplemental Figure S1). In 17 out of 18 litters, the same placenta was injected at each timepoint; in one dam (MNR, sham treated) fetal positioning over the placenta following the first treatment prevented subsequent treatments so the next closest placenta was injected for the second and third treatment. In the MNR + *IGF1* dams, placentas were separated based on receiving a direct injection of nanoparticle-mediated *IGF1* or by being indirectly exposed to circulating residual nanoparticle-mediated *IGF1* which was confirmed by expression levels of *hIGF1* via qPCR. Dams were sacrificed at GD60 (±3) via carbon dioxide asphyxiation followed by cardiac puncture and exsanguination. Fetuses (Control: n=8 female and n=11 male, MNR: n=5 female and 11 male, MNR + *IGF1*: n=6 female and 10 male) and the delivered placenta (placenta, sub-placenta, and decidua) were removed from the uterus and weighed. Fetal sex was determined at this time by examination of the gonads. Blood was collected via cardiac puncture from dams and fetuses at time of sacrifice. Fetal and maternal organs (heart, lung, liver, kidney, spleen, brain (fetus only), were isolated, weighed, and processed for histology (fixed in 4% PFA) or molecular analysis (preserved in RNAlater and snap-frozen). In order to reduce bias, all subsequent analyses were performed blinded to diet and nanoparticle treatment.

### RT-PCR and Quantitative PCR (qPCR)

Approximately 150–200 mg of placenta tissue was lysed and homogenized in RLT-lysis buffer (Qiagen) using a tissue homogenizer. RNA was extracted and DNase treated using the RNeasy Midi Plus Kit (Qiagen) following standard manufacturers protocols. The High-capacity cDNA Reverse Transcription kit (Applied Biosystems) was used to convert 1 µg RNA to cDNA, following standard manufacturers protocol. *IGF1* primers were designed for species specificity for both guinea pig (*gpIgf1; Fwd: 5’ –* CCTTCTGCTTGCTCGTCC, Rev: 5’ - TCTCCAGCCTCCGCAGGTCG) and plasmid-specific human *(hIGF1; Fwd: 5’-* CGCTGTGCCTGCTCACCT, Rev: 5’ – TCTCCAGCCTCCTTAGATCA) and purchased from Integrated DNA Technologies (IDT). Initially, gDNA and cDNA from guinea pig and human placenta were tested using RT-PCR to confirm specificity of both *gpIgf1* and *hIGF1*, respectively. Expression of plasmid-specific *hIGF1* <u>was then determined in guinea pig placenta using a RT-PCR protocol. In a 25 µL reaction, 10 µL of cDNA was mixed with</u> *<u>hIGF1</u>* <u>primers, positive control primers</u> *<u>RSP20</u>* <u>and</u> Fast Start PCR mastermix (Roche), following manufacturer’s instructions. RT-PCR was performed as follows: Activation, 95°C for 4 min; Denaturing-Annealing-Elongation, 95°C-60°C-72°C for 30s-30s-1min times 35 cycles; Final Elongation, 72°C for 7 min. RT-PCR samples were then run on a 1.5% agarose gel for 90 mins at 180V and imaged using a Chemidoc (Biorad) Imager (Supplemental Figure S2). To further explore differences in plasmid-specific *hIGF1* expression in the MNR + *IGF1* placentas that were directly injected or indirectly exposed, qPCR was then performed. qPCR reactions (20 µL containing 10 µL cDNA) were set up in duplicate and performed on the Quant 3 PCR System (Applied Biosystems) using PowerUp Sybr Green Master Mix (Applied Biosystems), following standard manufacturer’s instructions. Commercially available KiCqStart primers (Sigma) were used to measure mRNA expression of guinea pig *Igf2, Igf1 Receptor* (*Igf1R*) and *Igf Binding Protein 3* (*IgfBP3*), normalized to *β-actin, Gapdh* and *Rsp20*. Relative mRNA expression was calculated using the comparative CT method with the Design and Analysis Software v2.6.0 (Applied Biosystems).

### Serum/plasma analysis

Maternal (heparinized, EDTA and serum) and fetal (heparinized) blood samples were collected at time of sacrifice and serum/plasma obtained by centrifugation. Plasma was analyzed for maternal and fetal glucose and lactate using the YSI model 2700 Glucose/Lactate Analyzer following standard protocols. Maternal plasma electrolytes (sodium and potassium) were analyzed using the Roche Electrolyte Analyzer 9180 following standard protocols. Cortisol and progesterone in maternal serum and cortisol in fetal plasma were analyzed using commercially available ELISA kits (Arbor Assays) following manufacturers specifications. For cortisol, assay sensitivity was 27.6 pg/mL, the limit of detection was 45.4 pg/mL and inter-assay variability of 7.2-10.9%. For progesterone assay, sensitivity was 47.9 pg/mL, the limit of detection was 52.9 pg/mL and inter-assay variability of 4.1-7.0%.

### Statistical Analysis

All statistical analyses were performed using SPSS Statistics 29 software. Female and male fetuses were analyzed separately, except for analysis of plasmid-specific *hIGF1* expression. In the sham treated Control and MNR groups, there was no effect of direct placental injection for any outcomes measured and was therefore removed as a main effect in these groups. Distribution assumptions were checked with a Q-Q-Plot. For maternal outcomes, statistical significance was determined using generalized linear modeling with gamma as the distribution and log as the link function. Diet and nanoparticle-mediated *IGF1* treatment were considered main effects, and gestational age and litter size designated covariates. For placental expression of plasmid-specific *hIGF1*, statistical significance was determined using generalized estimating equations with gamma as the distribution and log as the link function only on MNR + *IGF1* placentas. Dams were considered the subject, fetal sex and direct inject or indirect exposure considered main effects and gestational age and litter size as covariates. For all remaining fetal and placental outcomes, statistical significance was determined using generalized estimating equations with gamma as the distribution and log as the link function. Dams were considered the subject, diet and nanoparticle-mediated *IGF1* treatment treated as main effects and gestational age and litter size as covariates. In male fetuses, direct injection or indirect exposure of nanoparticle-mediated *IGF1* was also considered as a main effect. Statistical significance was considered at P≤0.05. For statistically significant results, a Bonferroni post hoc analysis was performed. Results are reported as estimated marginal means ± 95% confidence interval.

## RESULTS

### IGF signaling is regulated in a sexually dimorphic manner in both Fetal Growth Restriction and following nanoparticle-mediated trophoblast-specific *IGF1* treatment

Repeated nanoparticle-mediated *IGF1* treatments from mid-pregnancy to near term resulted in no maternal health complications, no fetal loss, and no placental hemorrhage. Average litter size was 3 (minimum 1 and maximum 5), with no difference between Control and MNR diets (mean ± SEM: Control 3.2 ± 0.31 vs. MNR 2.9 ± 0.23, P=0.503). No resorptions were recorded for any pregnancies. One dam within the MNR + *IGF1* treatment group was determined to have a singleton pregnancy and was excluded from the study as the physiology of monozygotic pregnancies and polyzygotic pregnancies in guinea pigs differ significantly ^35-37^. 13 out of 17 dams (76%: 4 Control and 9 MNR) became pregnant on the first mating attempt, whilst the remaining 4 (24%: 2 Control and 2 MNR) became pregnant on the second mating. There was no difference in the number of female or male fetuses between Control or MNR diets (Control female fetuses = 8, male fetuses = 11 and MNR female fetuses = 9 and male fetuses = 21; Pearson Chi-Square = 0.752, p-value = 0.386). 12 out of 17 (71%: 3 Control and 9 MNR) dams had a uterine horn with a single fetus whilst the opposite uterine horn contained two or more fetuses. 3 dams (1 Control and 2 MNR) had all pups within one uterine horn and no fetuses in the opposite uterine horn, and 2 dams (1 Control and 1 MNR) had more than one fetus in both uterine horns.

One fetus in each pregnancy was chosen to receive a direct nanoparticle-mediated *IGF1* or sham injection into its placenta to study effects of the full direct treatment on a single fetus as most human pregnancy only harbor one fetus. Other fetuses in the litter were used to investigate possible circulating indirect exposure. As fetal sex determination is not possible at time of injection, the sex of the fetus receiving the direct injection verses the indirectly exposed littermates was random. At time of sacrifice fetuses were sexed, and it was determined that, in this study, only the placentas of male fetuses had received a direct nanoparticle-mediated *IGF1* injection. Therefore in data pertaining to outcomes from female fetuses, there is no direct injected MNR + *IGF1* treatment category. Female and other male littermates were indirectly exposed to circulating nanoparticle.

Repeated nanoparticle treatments with the human *IGF1* gene resulted in the expression of *hIGF1* mRNA within directly injected and indirectly exposed placentas, and not sham injected placentas at time of sacrifice (Supplemental Figure S1). Analysis using the more sensitive qPCR protocol confirmed indirectly exposed placentas had less *hIGF1* expression than directly injected placentas (Figure 1A). Endogenous levels of *gpIgf1* were reduced in MNR placentas in males compared to controls (Figure 1B; p<0.001), but this reduction failed to reach significance in the MNR females. No significant difference was seen in endogenous *gpIgf1* expression in MNR + *IGF1* placentas compared to controls in either sex (Figure 1C).

**Figure 1.**
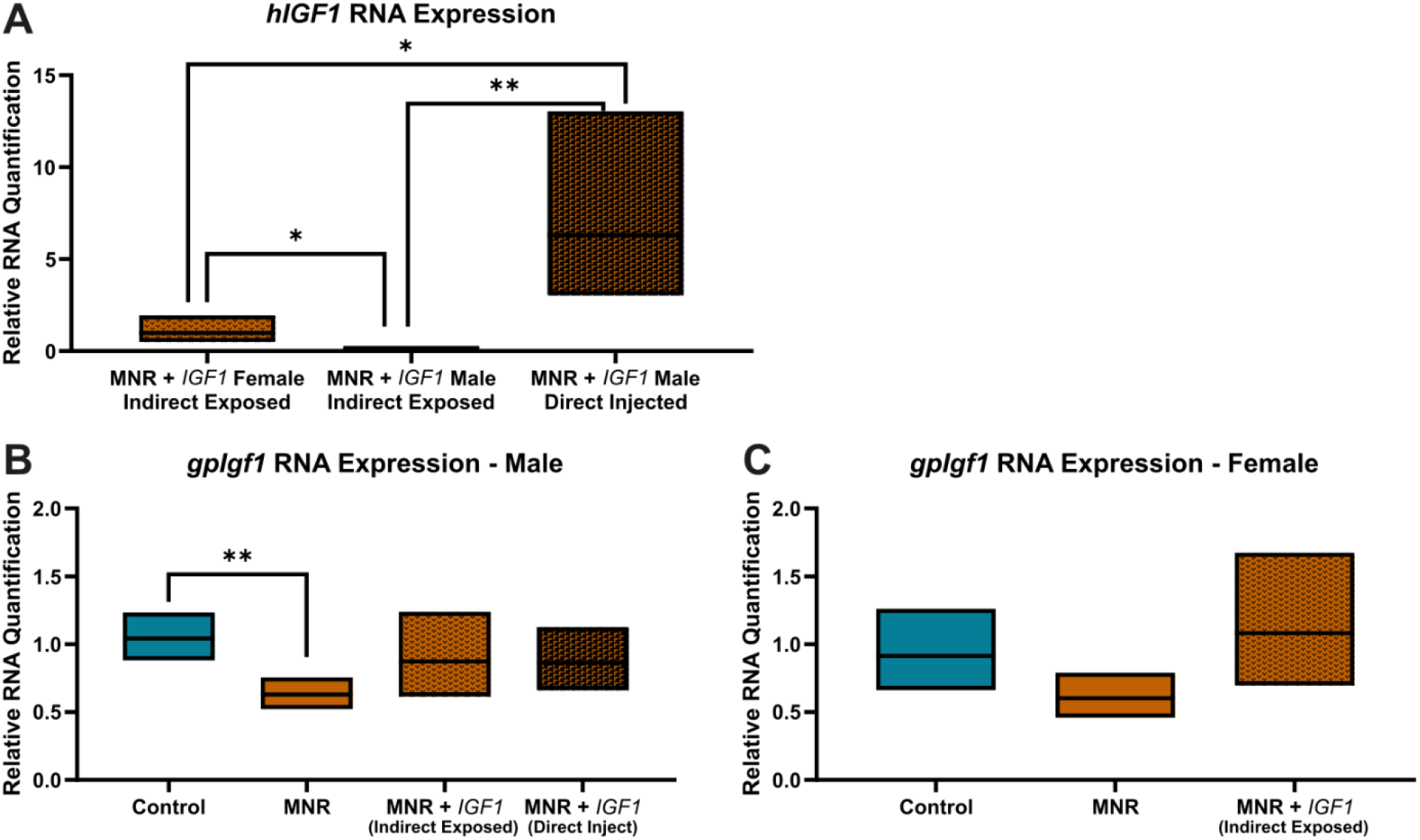
Effects of maternal nutrient restriction (MNR) and repeated nanoparticle-mediated *IGF1* delivery (MNR + *IGF1*) on plasmid-specific human *IGF1* (*hIGF1*) and endogenous guinea pig *Igf1* (*gpIgf1*). **A**. *hIGF1* mRNA was present within directly injected and indirectly exposed placentas from dams treated with the *hIGF1* nanoparticle, although indirectly exposed placentas had less. **B**. Endogenous *gpIgf1* was lower in the sham treated MNR placentas of male fetuses compared to sham treated Control; *gpIgf1* levels in the MNR + *IGF1* groups were comparable to Control. **C**. In placentas of female fetuses, there was no difference in endogenous *gpIgf1* between Control, MNR or MNR + *IGF1*. Control (+ sham treatment): n = 6 Dams (8 female and 11 male fetuses), MNR (+ sham treatment): n = 6 Dams (5 female and 11 male fetuses), MNR + *IGF1*: n = 5 Dams (6 female and 10 male fetuses). Data are estimated marginal means ± 95% confidence interval. P values calculated using generalized estimating equations with Bonferroni post hoc analysis. *P≤0.05; **P≤0.01

Sexually dimorphic changes also occurred in the placental expression of other IGF family members including *Igf2, Igf1R*, and *IgfBP3. Igf2* expression was not different from control in male MNR placentas (Figure 2A), but significantly increased in female MNR placentas (Figure 2B; P=0.037), indicating a potential compensatory response to maternal dietary restriction/stress by the female but not male placenta. Indirect exposure of males and females to nanoparticle-mediated *IGF1* did not further regulate *Igf2* expression compared to the MNR groups but direct *IGF1* treatment in the male placenta potentially increased *Igf2* expression (P=0.056). Indirectly exposed female placentas demonstrated increased *Igf1R* (Figure 2D; P=0.023) and *IgfBP3* (Figure 2F; P=0.02) expression in the MNR + *IGF1* group compared to controls while males showed no change (Figure 2C and 2E, respectively).

**Figure 2.**
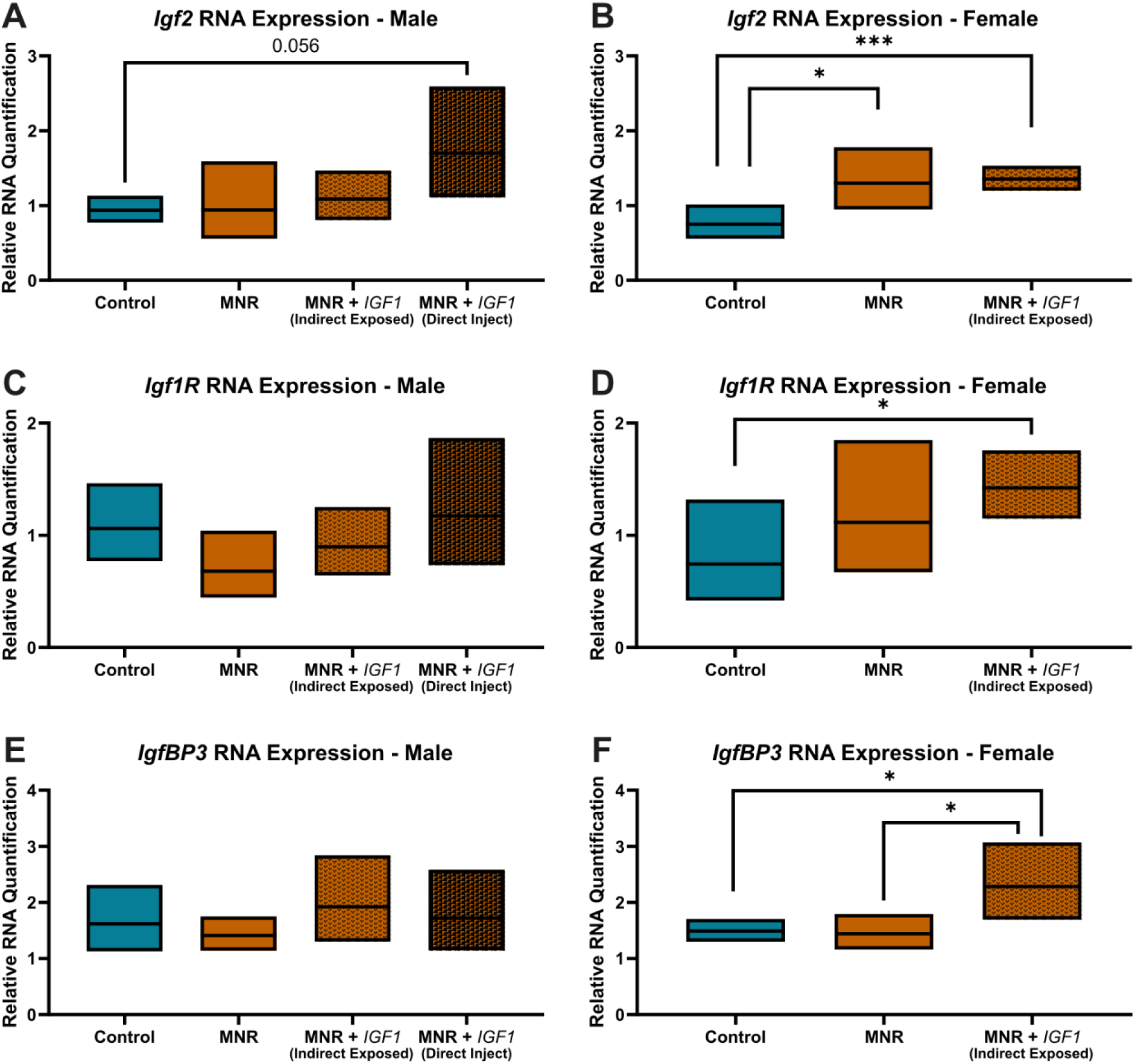
Effect of maternal nutrient restriction (MNR) diet, and repeated *IGF1* delivery on placenta *IGF1* signaling genes: **A**. *Insulin like growth factor 2* (*Igf2*) was unaltered among all male placental groups. **B**. Females in the MNR and MNR + *IGF1* groups had increased *Igf2* placental expression of compared to controls. **C**. *Insulin like growth factor 1 Receptor (Igf1R*) levels were unaltered between placentas of male fetuses. **D**. *Igf1R* expression was increased in MNR + *IGF1* indirectly exposed female placentas compared to controls. **E**. *Insulin like growth factor binding partner 3* (*IgfBP3*) was unaltered between placentas of male fetuses. **F**. *IgfBP3* expression was increased in MNR + *IGF1* indirectly exposed female placentas compared to control and MNR. Control (+ sham treatment): n = 6 dams (8 female and 11 male fetuses), MNR (+ sham treatment): n = 6 dams (5 female and 11 male fetuses), MNR + *IGF1*: n = 5 dams (6 female and 10 male fetuses). Data are estimated marginal means ± 95% confidence interval. P values calculated using generalized estimating equations with Bonferroni post hoc analysis. *P≤0.05; **P≤0.01, ***P≤0.001

### Repeated direct nanoparticle-mediated *IGF1* treatment ameliorates fetal growth restriction

Sham treated MNR fetuses of both sexes weighed significantly less than control fetuses at time of sacrifice (Males: Figure 3A; P=0.019, Females: Figure 3B; P=0.003). Fetuses whose placenta received direct nanoparticle-mediated *IGF1* injection (males only directly treated) weighed significantly more than MNR males (P=0.036) and the same as control males. Weights of male and female fetuses whose placentas were indirectly exposed to nanoparticle-mediated *IGF1* was increased 8% and 7%, compared to sham-treated MNR, demonstrating some recovery towards control weights but were not statistically significantly different from either Control or MNR. In males, Wald Chi-Square for direct injection of nanoparticle-mediated *IGF1* compared to indirect exposure was 7.51 (95% Wald CI: 0.025-0.148, P=0.006) indicating a significant association between level of *hIGF1* in the placenta and male fetal weight. Similarly, in female fetuses, Wald Chi-Square for nanoparticle-mediated *IGF1* treatment = 1.267 (95% Wald CI: -0.178-0.048) indicating a potential association with increased fetal weight.

**Figure 3.**
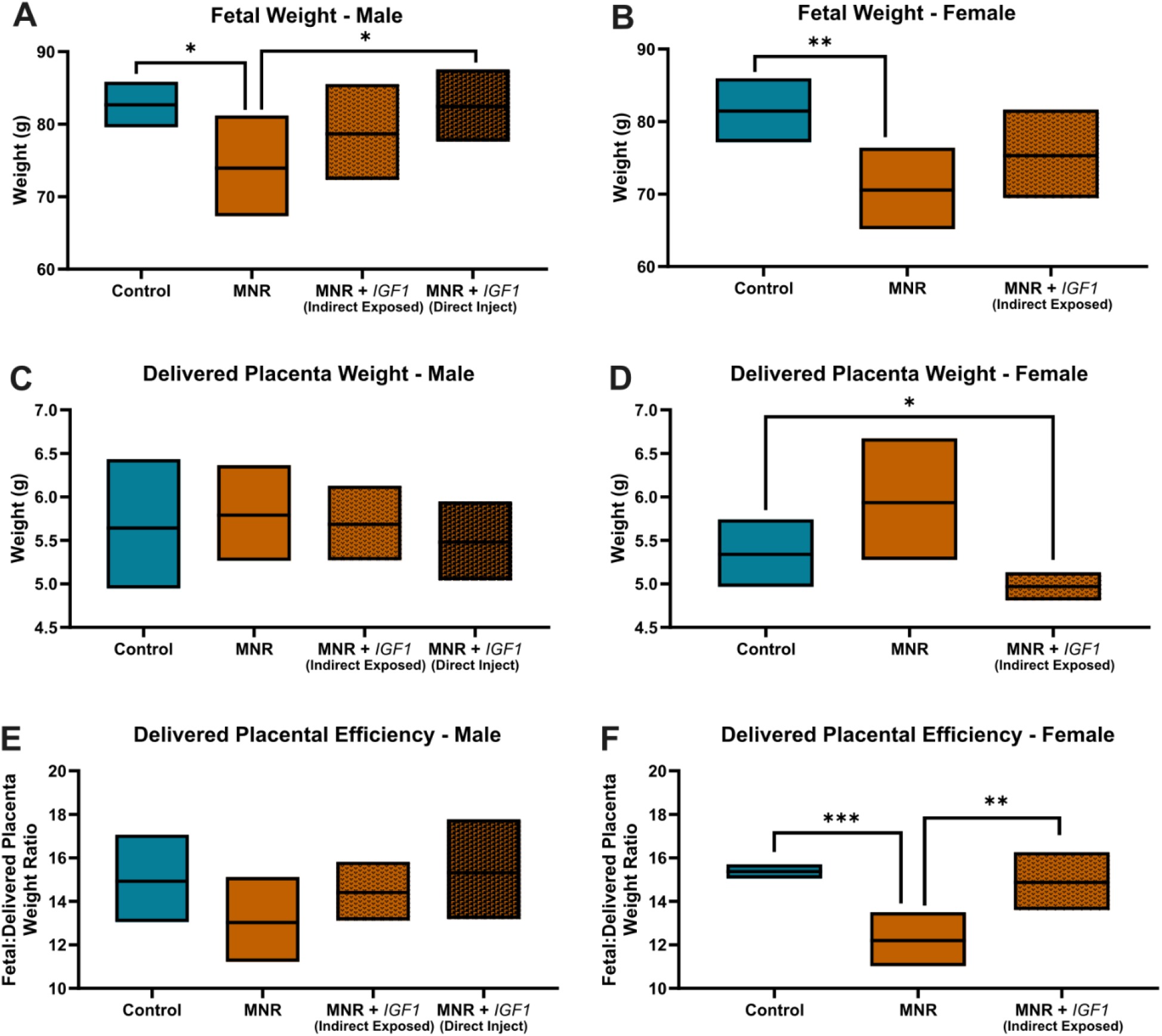
Effect of maternal nutrient restriction (MNR) diet, and repeated *IGF1* delivery on fetal and placental weight. **A & B**. MNR fetuses weighed significantly less than controls in both sexes. MNR + *IGF1* (direct injection and indirect exposed) fetuses showed comparable weights to controls in males and females with full correction from MNR with direct *IGF1* treatment in males. **C**. Delivered placenta weight showed no changes among any groups in male fetuses. **D**. Delivered placenta weight decreased in MNR + *IGF1* compared to controls in females. **E**. Delivered placental efficiency showed no changes among males in any group. **F**. Delivered placental efficiency decreased with MNR in females compared to controls but was corrected in the MNR + *IGF1* group. Control (+ sham treatment): n = 6 (8 female and 11 male), MNR (+ sham treatment): n = 6 (5 female and 11 male), MNR + *IGF1*: n = 5 (6 female and 10 male). Data are estimated marginal means ± 95% confidence interval. P values calculated using generalized estimating equations with Bonferroni post hoc analysis. *P≤0.05; **P≤0.01, ***P≤0.001

Males had no changes in delivered placenta weight between any of the groups (Figure 3C). In female fetuses, delivered placental weight was similar between MNR and control (Figure 3D) resulting in lower delivered placenta efficiency (fetal to delivered placental weight ratio) (Figure 3F, P<0.001). In the MNR + *IGF1* female fetuses, delivered placenta weight was lower compared to controls (Figure 3D. P=0.013). However, delivered placenta efficiency was similar between MNR + *IGF1* and controls, and increased compared to sham treated MNR (Figure 3F, P=0.011). Delivered placenta efficiency in male fetuses (Figure 3E) showed similar trends in changes with MNR and nanoparticle-mediated *IGF1* treatment which were not statistically different. However, Wald Chi-Square for nanoparticle-mediated *IGF1* treatment = 3.091 (95% Wald CI: -0.388-0.021) indicating a potential association with delivered placenta efficiency that may be more clearly determined with a larger sample size.

Fetal organs were dissected and weighed at time of necropsy. In the MNR group, male and female fetuses displayed brain sparing with significantly higher brain weights as a percentage of fetal weight compared to controls (Table 1). However, brain weight as a percentage of fetal weight was comparable to control in the MNR + *IGF1* directly treated male fetuses and the MNR + *IGF1* indirectly treated female fetuses. Additionally in male fetuses, liver weight as a percentage of fetal weight was increased in MNR (P=0.016) and in indirectly exposed MNR + *IGF1* (P<0.001) when compared to control, and lung weight as a percentage of fetal weight was increased in directly injected MNR + *IGF1* fetuses compared to control (P=0.002). In female fetuses, liver weight as a percentage of fetal weight was decreased in indirectly exposed MNR + *IGF1* fetuses compared to sham treated MNR (P=0.031). Lung weights as a percentage of fetal weight in female fetuses was comparable across groups.

**Table 1.**
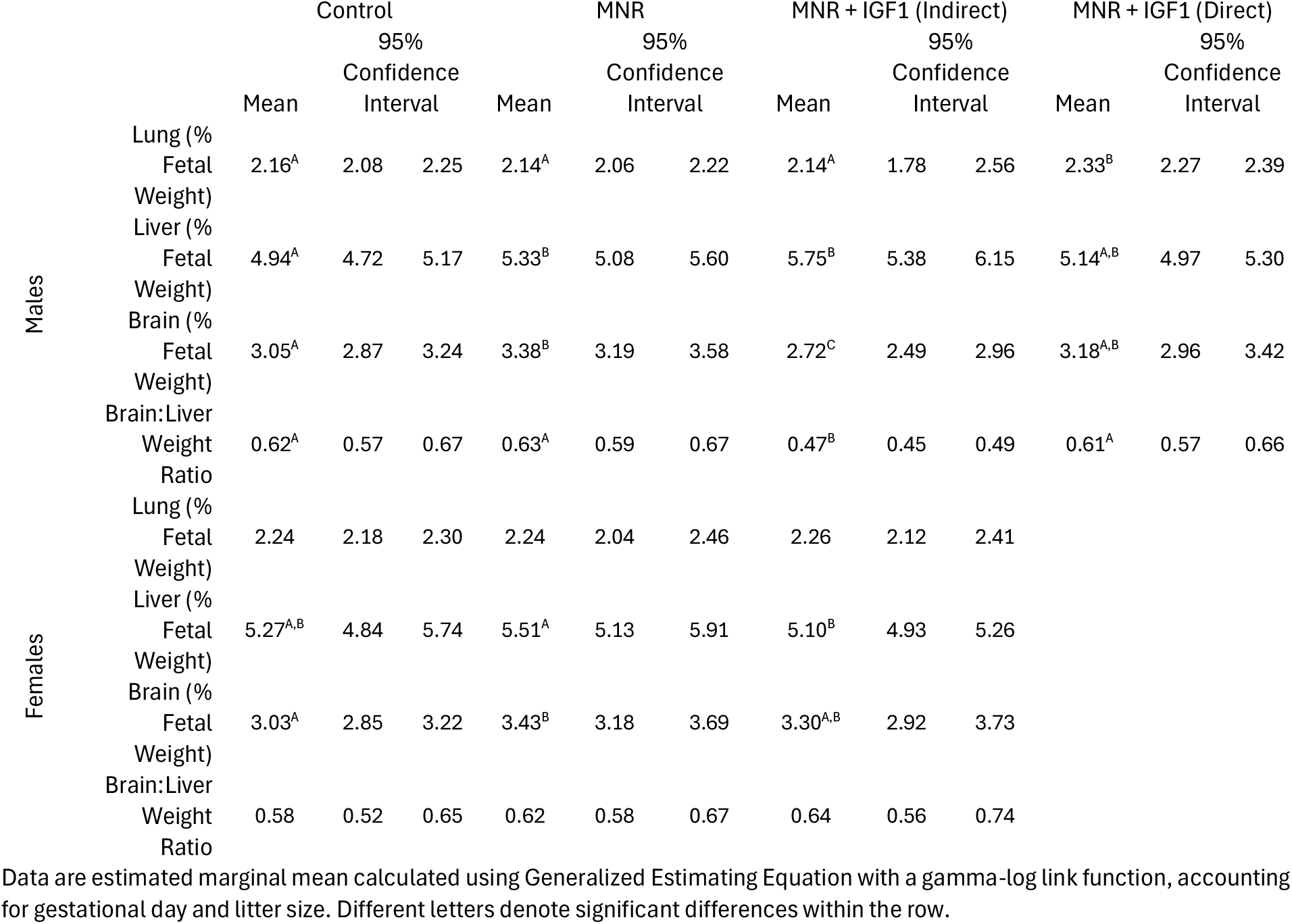
Fetal organ weights between control, MNR and nanoparticle-mediated *IGF1* treatment (MNR + *IGF1*) groups.

### Repeated nanoparticle-mediated *IGF1* treatment effectively corrects aberrant fetal glucose and cortisol levels

Glucose, lactate, and cortisol were measured in fetal blood at time of sacrifice. Male fetuses showed significantly reduced fetal blood glucose with MNR when compared to control (Figure 4A; P=0.001). However, MNR + *IGF1* fetuses, both direct and indirect, showed increased blood glucose when compared to MNR and comparable to control (Figure 4A; MNR + *IGF1* (Indirect) P = 0.019, MNR + *IGF1* (Direct) P = 0.039). Females, however, showed no changes in fetal blood glucose between groups (Figure 4B). In contrast, males showed no significant changes in blood lactate levels (Figure 4C), while females showed an increase blood lactate in the MNR and MNR + *IGF1* groups compared to controls (Figure 4D; MNR P=0.006, MNR + *IGF1* P<0.001). Blood cortisol levels were increased with MNR in both males (Figure 4E; P<0.001) and females (Figure 4F; P<0.001) when compared to control. However, blood cortisol levels were reduced with nanoparticle-mediated *IGF1* treatment as there was no difference between MNR + *IGF1* and control fetuses.

**Figure 4.**
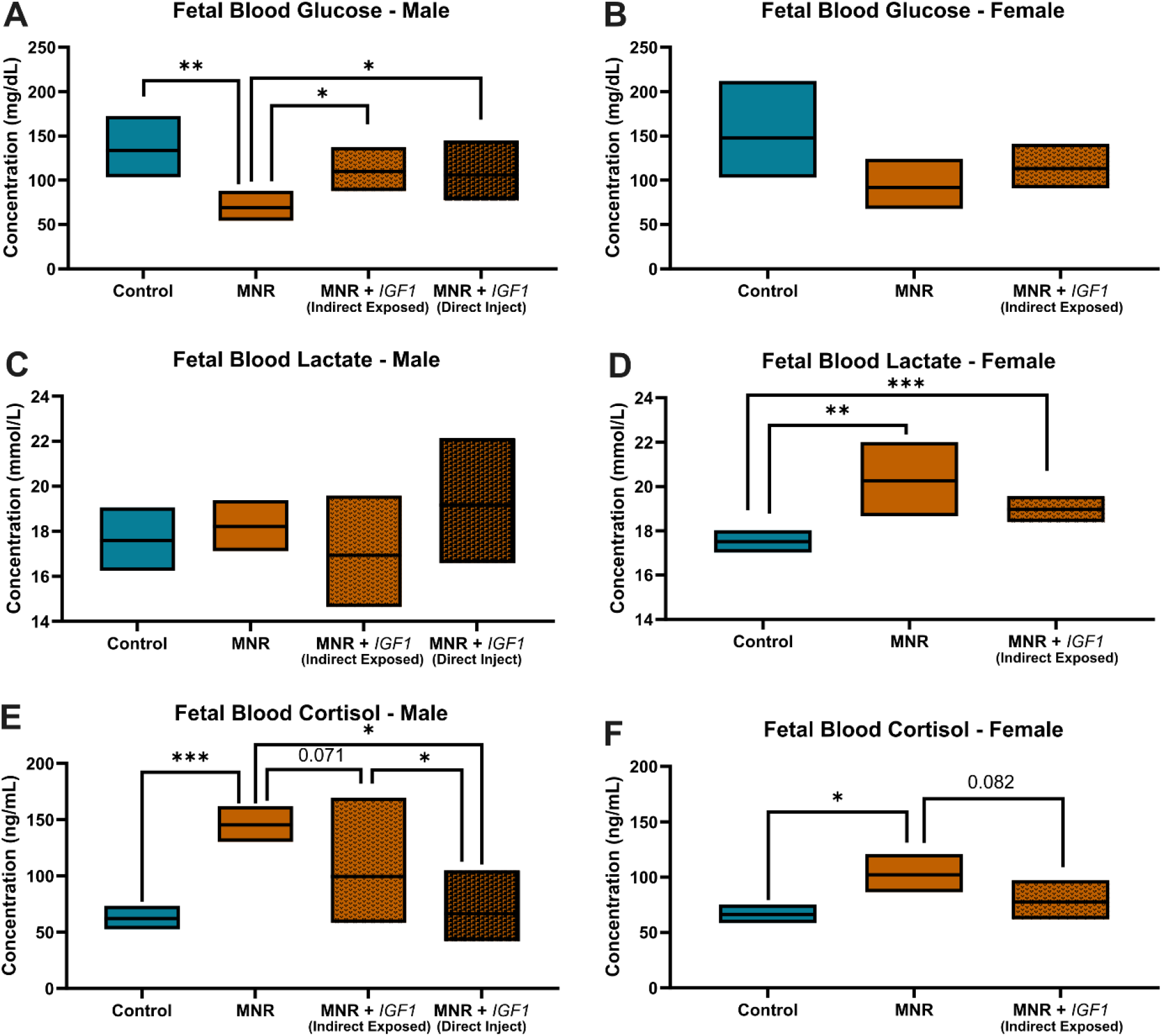
Effect of maternal nutrient restriction (MNR) diet, and repeated IGF1 delivery on fetal blood glucose, lactate, and cortisol. **A**. Male MNR fetuses had lower blood glucose levels compared to controls. MNR + *IGF1* direct injection and indirect exposure increased blood glucose back to control levels. **B**. There were no changes in female fetal blood glucose levels. **C**. Blood lactate levels were unaltered in male fetuses. **D**. Female fetuses had increased blood lactate with MNR and MNR + *IGF1*. **E & F**. There were increased blood cortisol levels with MNR in both males and females. MNR + *IGF1* male and female fetuses had blood cortisol levels comparable to control. Control (+ sham treatment): n = 6 dams (8 female and 11 male fetuses), MNR (+ sham treatment): n = 6 dams (5 female and 11 male fetuses), MNR + *IGF1*: n = 5 dams (6 female and 10 male fetuses). Data are estimated marginal means ± 95% confidence interval. P values calculated using generalized estimating equations with Bonferroni post hoc analysis. *P≤0.05; **P≤0.01, ***P≤0.001

### Maternal cortisol levels are elevated with MNR but return to control levels with repeated nanoparticle-mediated *IGF1* placental treatment

Dams had no significant changes in total carcass weight across groups, further indicating the establishment of FGR without using a model of starvation (Supplemental Table S1). Maternal organs were dissected and weighed. No differences were seen in these weights other than some changes in maternal liver weight shown in supplement (Supplemental Table S1). Maternal blood glucose, lactate, and cortisol levels were measured at time of sacrifice. Maternal blood glucose levels were unaltered between control, MNR and MNR + *IGF1* group, and remained within published physiological levels ^38^ (Figure 5A). Lactate, the end product of glucose metabolism and 2nd major fuel for fetal oxidative metabolism, was also unaltered between all groups (Figure 5B). Sodium and Potassium levels were also unchanged between all groups (Figure 5C an 5D, respectively). Maternal cortisol levels were significantly increased in the MNR group (P=0.011) but corrected to control levels with treatment in the MNR + *IGF1* group (P=0.013) (Figure 5E). Circulating maternal progesterone levels were unaltered between any of the groups (Figure 5F).

**Figure 5.**
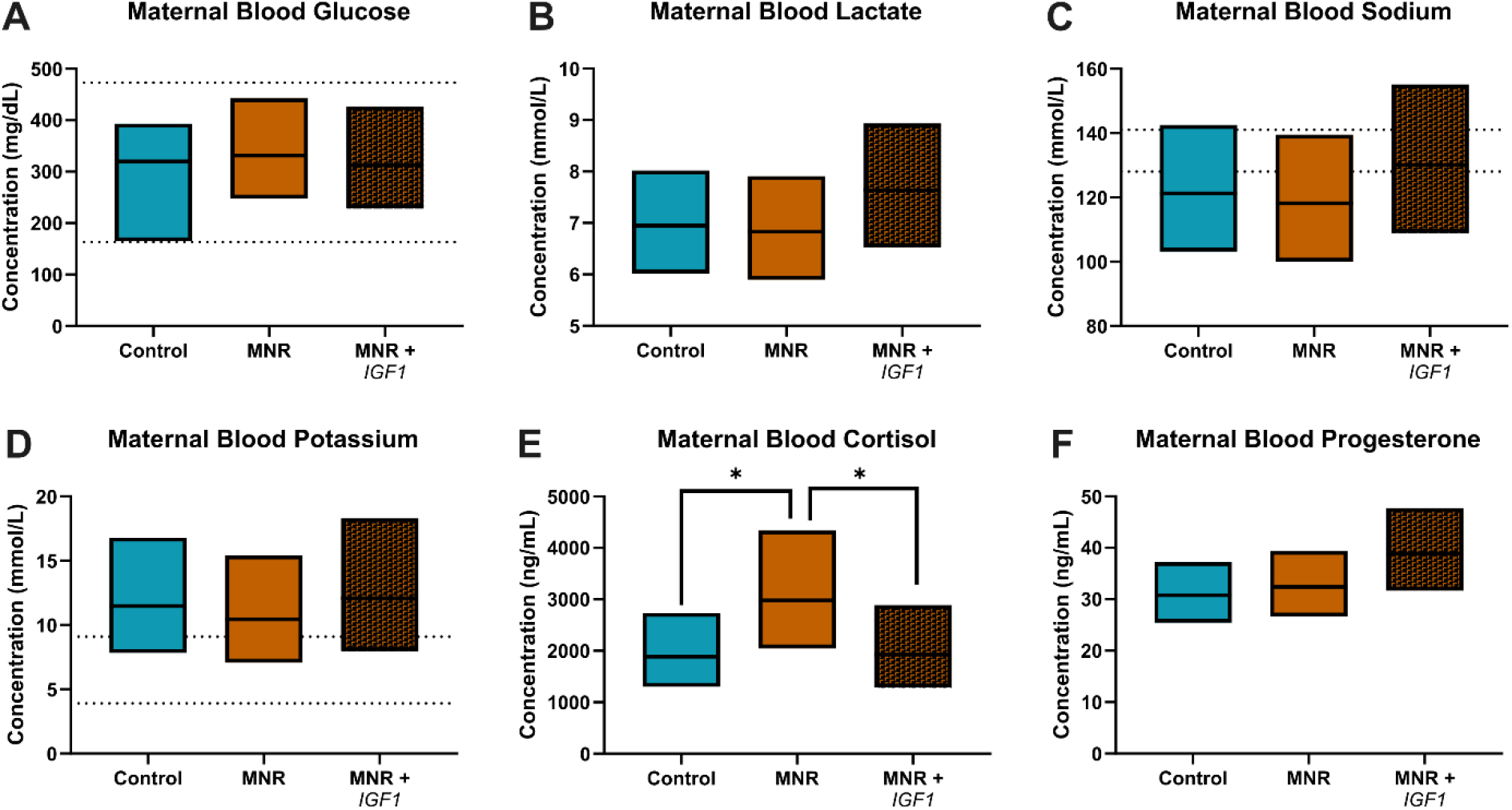
Effect of maternal nutrient restriction (MNR) diet, and repeated nanoparticle-mediated *IGF1* delivery on maternal blood biochemical measures: **A**. Maternal blood glucose was unaltered among groups **B**. Maternal blood lactate was unaltered among groups **C**. Maternal blood sodium was unaltered among groups **D**. Maternal blood potassium was unaltered among groups **E**. Cortisol was increased with MNR but corrected to control levels with MNR + *IGF1* **F**. Maternal blood progesterone was unaltered among groups. Dotted lines denote normal ranges for each measure in non-pregnant guinea pigs. Control (+ sham treatment) n=6, MNR (+ sham treatment) n=6, MNR + *IGF1* n=5. Data are estimated marginal means ± 95% confidence interval. P values calculated using generalized linear models with Bonferroni post hoc analysis. *P≤0.05; **P≤0.01, ***P≤0.001

## DISCUSSION

There are currently no treatments for fetal growth restriction nor placental maldevelopment/dysfunction. To establish a safe and effective treatment for placental insufficiency it is necessary to create a treatment that improves placental development, structure and/or function to increase fetal growth without 1. causing placentamegaly, 2. negatively impacting the maternal environment, or 3. causing negative side-effects to the fetus, or macrosomia in cases of misdiagnosed FGR. Placental *hIGF1* expression following nanoparticle-mediated *IGF1* treatment was associated with comparable fetal weight in male and female MNR + *IGF1* fetuses when compared to control, increased fetal blood glucose, and reduced blood cortisol levels in both fetuses and dams. Overall, this data, in addition to our previous studies on immediate effects of this *IGF1* gene therapy and the homeostatic mechanisms of the placenta when *IGF1* treatment is given to controls ^23^, show the potential of this therapy as a safe treatment for fetal growth restriction that positively impacts placental function, fetal growth, developmental programming, and improves the health of mother and offspring.

It is widely accepted that male fetuses are more susceptible to worse outcomes when affected by in utero disorders and perform more poorly in the NICU than female fetuses and neonates ^39, 40^. In Control fetuses, all fetal outcomes measured were comparable between females and males, however, in accordance with our previous investigations, we have identified sexually dimorphic responses with both MNR and our *IGF1* treatment in fetuses, recapitulating the human scenario ^24, 41, 42^. While we had no directly treated female placentas, indirectly treated female placentas demonstrated changes in IGF signaling components *Igf1R, IgfBP3* and *Igf2* mRNA whilst levels in both directly and indirectly treated placentas of males remained unchanged, highlighting dimorphic responses based on sex. We show that female placenta *Igf2* levels increase in response to maternal stress induced placental insufficiency potentially as a means to compensate for poor development and growth whereas male placentas do not. With the expression of *hIGF1* following indirect exposure combined with upregulation of *Igf2* from compensatory response, female placentas upregulate *Igf1R* and *IgfBP3* to likely regulate the bioavailability and response to increased *Igf1*. In contrast, male MNR placentas show no compensatory mechanism and only a trend towards increasing *Igf2* expression following direct *IGF1* treatment.

Both male and female fetuses, irrespective of whether they received direct injection or indirect exposure to *IGF1* treatment, showed improved growth with no statistical differences in fetal weights between controls and the *IGF1* treatment groups; importantly, males receiving direct treatment had rescued weight compared to the MNR group. This shows our novel treatment’s ability to effectively correct preexisting fetal growth restriction in a model of placental insufficiency. Placentas exposed indirectly to *IGF1* treatment showed an intermediary response between sham treated MNR and Controls and were not statistically different from either. In male fetuses, the indication of a dose response in fetal weight is clear between direct injection and indirect exposure and may still represent significant physiological improvement as any weight gain in utero is greatly valued in FGR fetuses. In addition to IGF family member regulation, delivered placenta weights and efficiency also showed sexually dimorphic responses. Whilst we cannot determine the impact of direct *IGF1* treatment to the female placenta from the current study, our data, in addition to historic reporting of females performing better in utero and in NICU under the same conditions as males, indicates the possibility that females may require a lower dosage compared to males for full correction of FGR ^39^.

In addition to improving placental signaling and fetal growth, repeated nanoparticle-mediated *IGF1* treatment impacted fetal brain growth. Asymmetric growth is a characteristic of 70-80% of all human FGR cases (and the MNR guinea pig model) with disproportionate growth causing increased blood supply to the fetal brain to maintain life and resulting in brain sparing effects ^43, 44^. This brain sparing maintains brain growth and leads to an increase in fetal brain weight as a percentage of total body weight compared to average for gestational age (AGA) or symmetrical FGR fetuses ^45, 46^. We found brain sparing effects in both male and female fetuses in our MNR group, but importantly comparable brain:body ratios in the *IGF1* treatment groups compared to controls, demonstrating that we are correcting full body weight and brain sparing. As a consequence of asymmetrical growth, organs such as the liver, kidneys, and spleen may be deprived of blood flow and growth and development are impaired. However, while some studies show reduction in these organ weights, others show no change in weight, but strong functional deficits ^12, 47, 48^. The abnormal growth and development of these organs from FGR are what lead to childhood and adulthood morbidities that increase risks associated diseases such as fatty liver disease, obesity, and cardiovascular disease ^6, 7, 33^. Studies identifying the precise malperfusions and functional consequences have been and continue to be investigated for these various organ systems ^49-54^. We show that a non-viral, polymeric placenta gene therapy treatment can improve fetal growth trajectories and one of the most prominent and significantly disordered features of FGR. With correction of brain sparing and increased growth in some of these peripheral organs with our *IGF1* therapy we may be able to reduce the probability of developing many cognitive and peripheral organ morbidities later in life.

Appropriate fetal development and growth is dependent on the transfer of oxygen and nutrients from the maternal to fetal circulation. Blood glucose and oxygen are reduced in FGR fetuses often as a result of aberrant placenta structure and/or function, impeding oxygen diffusion or failing to transport enough nutrients into fetal circulation ^55-57^. Glucose is the main source of energy for almost every tissue type in the body ^58^. Fetal metabolism and growth will impact an individual’s body functions and energy metabolism for the rest of their lives, so if a fetus is receiving less than adequate levels of energy then all the organ systems of the body are put under stress. In our model, MNR male fetal blood glucose is significantly reduced compared to controls. Our treatment, however, corrects these decreased glucose levels with both direct injection and indirect exposure to *IGF1* treatment to the male placentas. Blood glucose was not statistically changed with MNR or nanoparticle-mediated *IGF1* treatment in females, possibly due to the females having less changes in endogenous *gpIgf1* levels as well as their compensatory *Igf2* signaling changes with MNR and therefore not leading to significant changes in glucose transport. Lactate is the end product of anaerobic metabolism of glucose and many studies have shown that it is produced in the placenta and delivered to both the maternal and fetal circulations. This lactate is used for oxidation, and increased levels of lactate have been correlated with fetal oxygenation and FGR ^59^. FGR fetuses have higher lactate levels than AGA fetuses as seen in our female MNR fetuses ^60^. Indirect exposure of the female placentas was not able to reduce these elevated fetal blood lactate levels. However, this may be a consequence of there being less *hIGF1* expressed in the placentas of female as they only indirectly received the *IGF1* nanoparticle treatment, and reduced blood lactate levels may be observed if placental expression of *hIGF1* was greater. In this study we demonstrate changes in blood glucose and lactate levels with placental insufficiency, and correction of blood glucose levels with our nanoparticle-mediated *IGF1* treatment.

It is well established that increased cortisol in pregnancy can cause disruption to fetal organ development and metabolism that leads to increased disease risk later in life, a process described as fetal programming (to which the entire DOHaD field is dedicated to understanding these mechanisms) ^61-65^. In MNR fetuses of both sexes we identified statistically significant increases in cortisol levels. This elevation is likely due to a combination of the fetus receiving less than adequate levels of nutrients for proper growth and reduced handling of maternal cortisol in the insufficient FGR placentas as seen in human cases that leads to increased cortisol crossing the placenta and impacting fetal organ development ^65-68^. Several studies have found increased maternal and fetal cortisol levels during pregnancy lead to increased risk of cognitive delays in the offspring, metabolic diseases, and cardiovascular disorders ^61, 69, 70^. Here, we induce biologically relevant reductions of cortisol in both fetuses whose placentas were indirectly exposed, and complete reduction to control levels in fetuses whose placentas received direct nanoparticle-mediated *IGF1* treatment. This correction of fetal cortisol shows immense potential for creating a better pregnancy environment that could lead to reduced risks for countless multigenerational adult morbidities creating healthier humans and vastly decreasing medical care needs and expenses.

It is equally important to ascertain the impact of this treatment on maternal physiology. Maternal blood measures included in this study; glucose, lactate, potassium, sodium and progesterone, represent commonly and easily measurable biochemical assays throughout human pregnancy. Our aim was to understand the impact of our treatment, if any, on the maternal system as aberrant changes in these levels could indicate negative impact on maternal physiology. We showed that glucose, lactate, sodium, potassium, and progesterone levels remained unchanged across all groups. Unsurprisingly, as our MNR model and human FGR have been linked to increased maternal stress, the maternal cortisol levels were elevated. In humans, elevated maternal cortisol not only may cross the insufficient placenta but also cause vasoconstriction of the uterine arteries leading to reduced blood flow in the placenta and poor fetal growth ^71^. Furthermore, maternal cortisol increase with FGR in humans has been linked to higher risks of developing postpartum depression, coronary heart disease, and post-traumatic stress disorder ^65, 72^. This is the first study to demonstrate that with treatment of the placenta, however, full correction of maternal cortisol levels back to control levels. While more understanding of the underlying mechanism is needed to fully understand these changes, the dams are still on a restricted diet which indicates reduced maternal cortisol is likely due to improved placental handling of the cortisol and restored placental signaling to the maternal system. This report shows that a non-viral, polymeric nanoparticle placental gene therapy for the treatment of FGR is capable of improving the fetal, placental, and maternal environments.

A limitation of this study was the lack of female fetuses receiving direct injection of our nanoparticle-mediated *IGF1*. We have previously shown that all placentas in the litter express the plasmid-specific *hIGF1* following a single nanoparticle-mediated *IGF1* injection into one placenta ^24^, and chose to continue this approach. However, fetal sex cannot reliably be determined via ultrasound at mid-pregnancy and prior to our gene therapy delivery, hence we cannot control for which sex gets directly treated. Studies in non-human primates which carry a singleton pregnancy are underway to assess translational aspects including dose, delivery and safety and will address both fetal sexes. Further mechanistic studies in the guinea pig into precise sexual dimorphic responses to the nanoparticle-mediated *IGF1* will also be conducted as sex as a biological variable is a pivotal factor in all aspects of fetal and placental development.

In the present study, we show the ability of our novel nanoparticle-mediated *IGF1* gene therapy to restore fetal growth to levels comparable with control with repeated treatment in a guinea pig model. Positive changes to fetal growth trajectories were associated with increased fetal blood glucose and reduced fetal cortisol, indicative of improved fetal physiology. Additionally, we showed no change in any maternal physiological measures while also correcting increased blood cortisol levels. We show a non-viral, polymeric gene therapy that increased placental *IGF1* expression can improve the entire pregnancy environment: maternal, placental, and fetal. By improving each of these environments we may be able to help improve the health of mothers and offspring through the rest of their lives. This combined with our previous studies using this therapy at both mid pregnancy and in numerous cell and animal models demonstrate the plausibility of this therapy for future human translation. By correcting fetal growth restriction and asymmetric growth we have the potential to prevent countless stillbirths, reduce preterm birth rates, decrease long NICU stays, and decrease numerous morbidities associated with the developmental origins of disease for overall healthier lives.

## Data availability

All data needed to evaluate the conclusions in the paper are present in the paper and/or the Supplementary Materials.

## Acknowledgments

We would like to thank Drs Craig Duvall and Mukesh Gupta for providing the co-polymer, veterinary and technical staff at the University of Florida Animal Care Services, and Khanh Huynh, Aditya Mahadevan, Dr Jason Puglise, and Dr Erica Smith for their assistance with animal necropsies.

## Author Contributions

BND performed experiments, analyzed data, and wrote manuscript. RLW conceived the study, performed experiments, analyzed data, and wrote manuscript. AW assisted with animals and performed experiments. HNJ obtained funding, conceived the study, and edited the manuscript. All authors approve the final version of the manuscript.

## Funding

This study was funded by Eunice Kennedy Shriver National Institute of Child Health and Human Development (NICHD) award R01HD090657 (HNJ) and K99HD109458 (RLW).

## Competing Interests

The authors have declared that no competing interest exists

## Ethics approval

Animal care and usage was approved by the University of Florida Intuitional Animal Care and Usage Committee (Protocol #202011236).

## Notes

### Competing Interest Statement

The authors have declared no competing interest.

### Summary of Updates

Manuscript revised based on peer-review. Extended explanation of methods. Additional results. Figures updated. Supplemental material updated.

