## Supplemental Material for "Placental Nanoparticle-mediated IGF1 Gene Therapy Corrects Fetal Growth Restriction in a Guinea Pig Model"

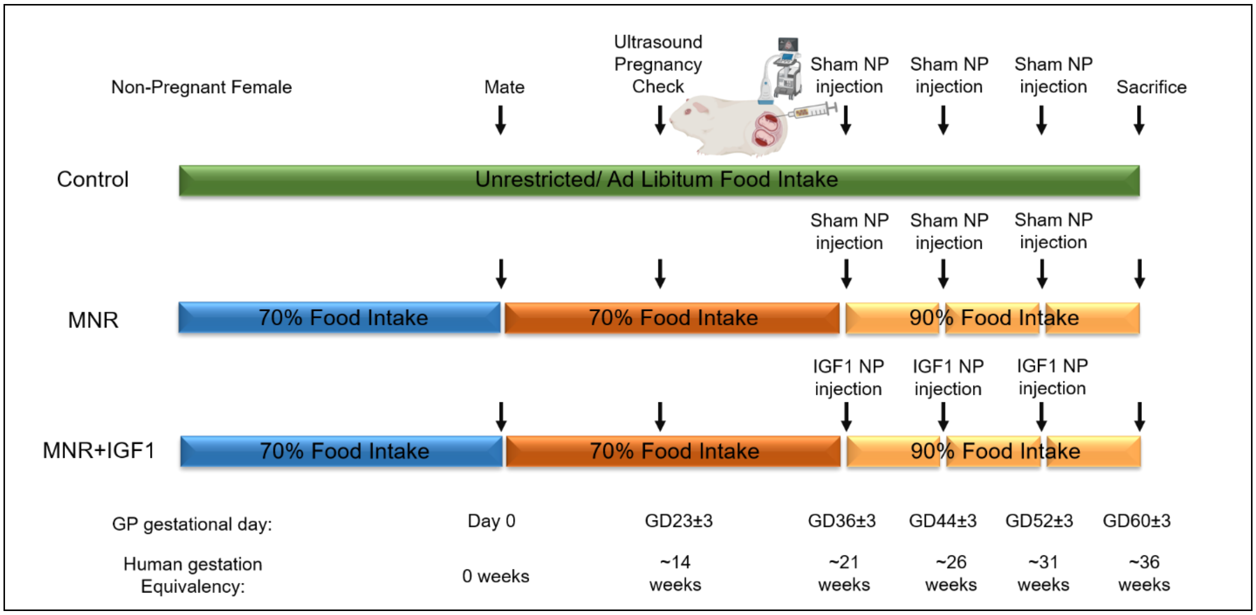


**Supplemental Figure S1.** Schematic of study design

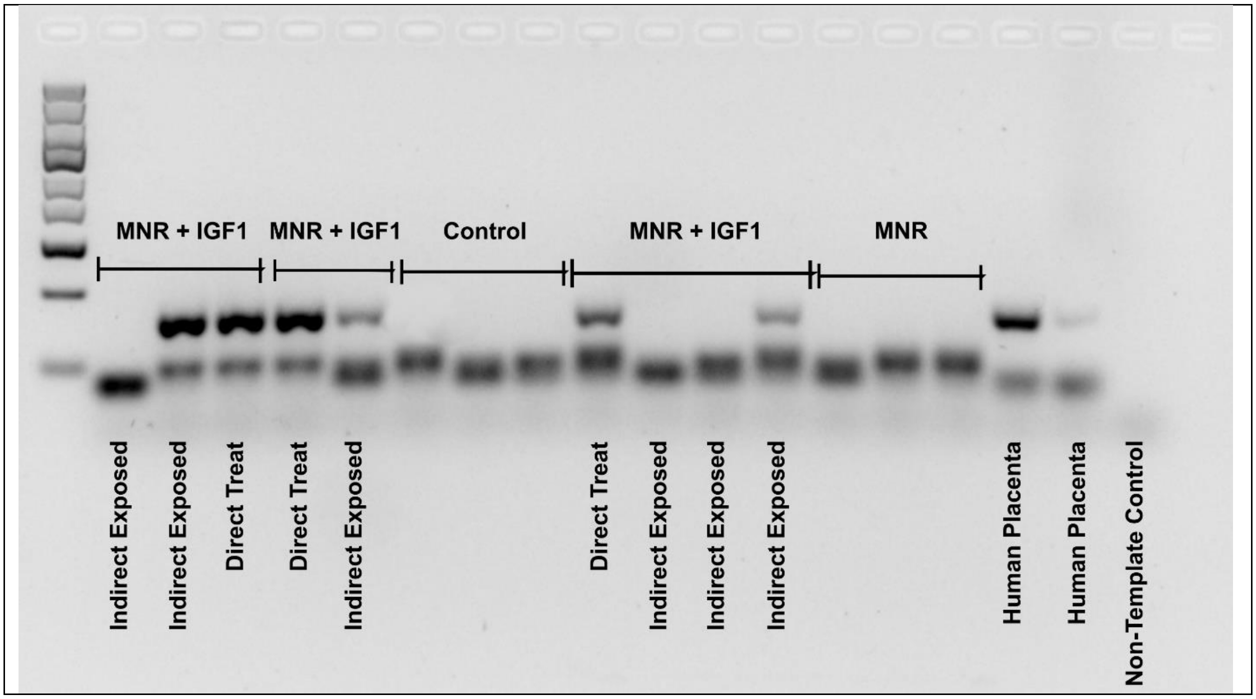


**Supplemental Figure S2.** Representative image of RT-PCR results confirming species specificity of *hIGF1* primers, and *hIGF1* expression in nanoparticle-mediated *IGF1* treated placentas. *hIGF1* (upper band) was present in directly injected MNR + *IGF1* placentas, and some indirectly exposed MNR + *IGF1* placentas. *hIGF1* was not present in any sham treated Control or MNR placentas. *RSP20* (lower band) was present in reactions from all samples. Human placenta samples were included as positive controls.

**Supplemental Table S1.** Maternal organ weights between control, MNR and nanoparticle-mediated *IGF1* treatment (MNR + *IGF1*) groups

| | Control | | | MNR | | | MNR + *IGF1* | | |
|---|---|---|---|---|---|---|---|---|---|
| | Mean | 95% Confidence Interval | | Mean | 95% Confidence Interval | | Mean | 95% Confidence Interval | |
| Carcass Weight (g) | 778 | 724 | 837 | 865 | 802 | 932 | 732 | 676 | 793 |
| Heart (% Carcass Weight) | 0.41 | 0.36 | 0.46 | 0.37 | 0.33 | 0.42 | 0.42 | 0.36 | 0.48 |
| Lung (% Carcass Weight) | 0.71 | 0.63 | 0.80 | 0.65 | 0.57 | 0.73 | 0.71 | 0.62 | 0.82 |
| Liver (% Carcass Weight) | 3.15[A] | 2.90 | 3.41 | 3.67[B] | 3.39 | 3.99 | 3.52[B] | 3.23 | 3.84 |
| Kidney (% Carcass Weight) | 0.73 | 0.69 | 0.78 | 0.75 | 0.70 | 0.80 | 0.75 | 0.71 | 0.81 |
| Spleen (% Carcass Weight) | 0.12 | 0.10 | 0.13 | 0.10 | 0.09 | 0.12 | 0.12 | 0.12 | 0.14 |

Data are estimated marginal mean calculated using Generalized Linear Model with a gamma-log link function, accounting for gestational day and litter size. Different letters denote significant differences between the row.
